# Synaptic control of temporal processing in the *Drosophila* olfactory system

**DOI:** 10.1101/2021.05.03.442428

**Authors:** David Fox, Katherine I. Nagel

## Abstract

Temporal filtering of sensory stimuli is a key neural computation, but the way such filters are implemented within the brain is unclear. One potential mechanism for implementing temporal filters is short-term synaptic plasticity, which is governed in part by the expression of pre-synaptic proteins that position synaptic vesicles at different distances to calcium channels. Here we leveraged the *Drosophila* olfactory system to directly test the hypothesis that short-term synaptic plasticity shapes temporal filtering of sensory stimuli. We used optogenetic activation to drive olfactory receptor neuron (ORN) activity with high temporal precision and knocked down the presynaptic priming factor unc13A specifically in ORNs. We found that this manipulation specifically decreases and delays transmission of high frequencies, leading to poorer encoding of distant plume filaments. We replicate this effect using a previously-developed model of transmission at this synapse, which features two components with different depression kinetics. Finally, we show that upwind running, a key component of odor source localization, is preferentially driven by high-frequency stimulus fluctuations, and this response is reduced by unc13A knock-down in ORNs. Our work links the extraction of particular temporal features of a sensory stimulus to the expression of particular presynaptic molecules.

## Introduction

Many behaviors depend on the ability of the brain to extract temporal information from complex sensory signals. Visual motion is detected by comparing signals that are displaced in space and differentially filtered in time (Behnia et al. 2014; Gruntman et al. 2019; Kim et al. 2014; Fransen and Borghuis, 2017; Hanson et al. 2019). Recognition of conspecific calls in frogs and birds depends on the selectivity of central neurons for patterns of acoustic stimulation over time (Edwards et al. 2002, Elliot et al. 2011, Schneider and Woolley 2013). Cerebellum-like circuits generate temporally diverse representations of motor commands (Kennedy et al. 2014) to facilitate motor learning (Suvrathan et al. 2016). In *Drosophila* and *C. elegans*, olfactory navigation depends on sensitivity to temporal changes in odor concentration (Gorur-Shandilya, et al. 2017; Lockery, 2011; Schulze et al. 2015; Nagel and Wilson 2011, Demir et al. 2020). While neurons with different temporal filtering properties are found throughout the brain, the biophysical mechanisms that support these different forms of temporal filtering are less clear.

One mechanism for controlling the temporal information extracted by central neurons is short-term synaptic plasticity. Synapses are regulated by past activity on timescales of milliseconds to seconds, collectively termed short-term plasticity (STP). Computational modeling suggests that different forms of STP can shape temporal filtering (Abbott et al. 1997; Abbott and Regehr 2004). At a molecular level, STP often depends on the spatial organization of presynaptic vesicle pools and their relationship to voltage-gated calcium channels (Zucker and Regehr, 2002), which can be differentially tuned in different synapse types (Pouille and Scanziani, 2004; Stokes and Isaacson 2010; Fulterer et al. 2018, Pooryasin et al 2020). While the molecular basis of STP in slice preparations has been well-studied, the functional role of synaptic molecular composition in sensory processing and animal behavior has been challenging to investigate experimentally.

In this study, we set out to test the hypothesis that the molecular makeup of synapses shapes temporal filtering of sensory information, with consequences for behavior. We focused on the *Drosophila* olfactory system, because it allows us to precisely manipulate the expression of molecular factors at a particular synapse and to monitor the timecourse of both neural activity and behavior. Due to turbulence, natural odor stimuli are broken into intermittent filaments that fluctuate across a range of frequencies (Celani et al. 2014; Connor et al. 2018.). Insect navigation in such stimuli has been shown to depend on the temporal characteristics of the stimulus (Baker et al. 1985; Mafra-Neto and Carde 1994, Alvarez-Salvado et al. 2018, Demir et al. 2020). The synapse between first and second-order olfactory neurons, known as olfactory receptor neurons (ORNs), and olfactory projection neurons (PNs), exhibits short-term depression (Kazama and Wilson, 2008, Nagel et al. 2015, Martelli and Fiala 2019). Computational modeling suggests that this depression contributes to its frequency filtering and nonlinear gain (Kazama and Wilson, 2008, Nagel et al. 2015). Recent work showed that knock-down of the priming factor unc13A, which helps to position synaptic vesicles at nanometer distances from voltage-gated calcium channels, specifically reduces a fast, rapidly-depressing component of release at ORN-PN synapses and can influence olfactory behavior (Fulterer et al. 2018, Pooryasin et al 2021). However, how this change in STP translates into altered temporal coding in the intact circuit is not clear.

To investigate the role of STP on sensory processing, we combined molecular manipulation of ORN-to-PN transmission with optogenetic activation of ORNs, and recorded the activity of both ORNs and PNs to well-controlled light stimuli. We found that unc13A was required specifically for transmitting high-frequency ORN firing rate fluctuations with short phase lag, implying that the molecular composition of the presynaptic terminal can shape temporal filtering of sensory information. We show that unc13A amplifies the response to distant filaments in a temporally complex plume. Finally we show that these effects of unc13A knock-down can be recapitulated using a previously developed two-component model of ORN-PN synaptic transmission (Nagel et al., 2015).

Second, we investigated whether STP at the ORN-to-PN synapse influences the dynamics of olfactory navigation behavior. Consistent with recent studies (Demir et al. 2020), we found that high frequency stimulus fluctuations promoted stronger and more consistent upwind runs, a key component of olfactory navigation. Knock-down of unc13A reduced this response to high frequency stimulus fluctuations, indicating that altered temporal filtering at ORN-PN synapses has consequences for behavioral feature extraction. Because many behaviors rely on the extraction of temporal information from ongoing stimulus fluctuations, and because short-term synaptic depression is found in many circuits, this mechanism should have a broad relevance for how sensory processing and behavior are set by changing the molecular properties of individual synapses.

## Results

### Unc13A promotes larger and faster responses to high frequency fluctuations in ORN firing rate

We first examined the effects of unc13A knock-down on temporal filtering. In a previous study we showed that ORN-PN synapses can transmit a wide range of temporal frequencies, in part because they have two components of transmission— a fast, rapidly adapting component and a slower, slowly-adapting component (Nagel et al. 2015). We therefore hypothesized that knock down of unc13A, which is required for fast rapidly-adapting synaptic transmission (Fulterer et al. 2018), would reduce high frequency transmission.

To test this hypothesis, we expressed Chrimson in ORNs using the orco promoter and presented forward and reverse frequency sweeps of red light intensity (628 nm) from 0.1 to 5 Hz at 60*μ*W/cm^2^ (Fig. 1A). We recorded both ORN and PN responses to the same light stimuli in control animals, and in animals in which unc13A was knocked down in orco+ ORNs using RNAi. We previously showed that this manipulation does not affect ORN spiking responses (Fulterer et al. 2018).

**Figure 1.**
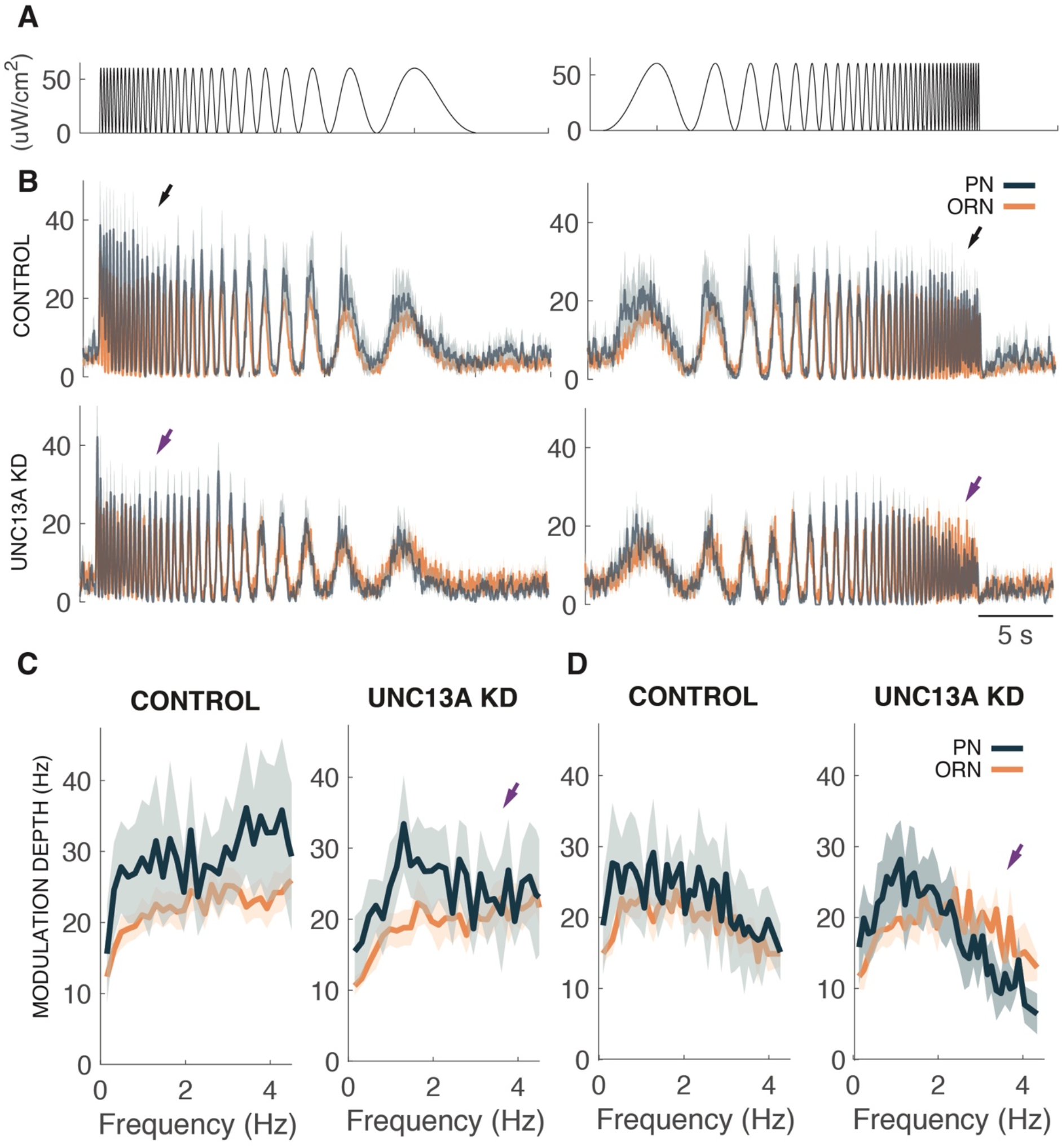
Knockdown of unc13A at ORN-PN synapses attenuates transmission of high frequency stimulus fluctuations. (A) Stimulus timecourses: frequency sweep stimuli spanning a frequency of 0.1–5 Hz and a luminance range of 0 to 60 uW/cm^2^ were presented in both reverse (high frequency first) and forward (low frequency first) directions. (B) Average firing-rate responses of ORNs (orange) and PNs (blue) to frequency sweep stimuli in (A). Top: Control responses (mean +/– SEM of resampled jackknife traces, n = 13 ORNs, n = 9 PNs. Same cells as in Fig. 1). Bottom: unc13A KD responses (n = 15 ORNs, n = 8 PNs). Arrows indicate where responses in unc13A KD are attenuated relative to control. (C) Modulation depth of PN and ORN firing rates as a function of stimulus cycle frequency for the reverse frequency sweep in control (black) and unc13A KD (violet). Arrow indicates decrease in PN response to high frequencies in unc13A KD (arrow). (D) Same as C for the forward sweep.

In control animals, we found that PN responses to the reverse sweep were larger at high frequencies than at low frequencies, while responses to the forward sweep were similar across frequencies (Fig. 1B–D). The larger responses at high frequencies likely reflect both short-term temporal filtering and slow time-scale adaptation, as the high frequencies occur at the beginning of the reverse chirp stimulus. In unc13A KD flies, no amplification of high frequency responses was observed for the reverse chirp, and responses to high frequencies were smaller than ORN input for the forward chirp. Thus, knock-down of unc13A reduced PN responses to high frequency ORN firing rate fluctuations regardless of stimulus direction (Fig. 1C–D).

We next examined the detailed phase relationship between ORN and PN firing rate responses. For the reverse frequency sweep, we observed that both ORN and PN responses peaked before the stimulus at low frequencies, consistent with the idea that PNs respond to the rate of change in the stimulus in this regime (Fig. 2A, left). In contrast, at high frequencies the peak of ORN responses occurred after the stimulus peak, and PN responses lagged ORN responses (Fig. 2A, right). In unc13A KD flies, the time lag of PN responses at high frequencies was more severely delayed, while there was no significant delay in PN responses at low frequencies (Fig. 2B–D). We did not observe such a phase delay with the forward sweep, perhaps because the contribution of unc13A is reduced at the end of the stimulus following synaptic depression. Thus, unc13A contributes to short latency PN responses especially when high stimulus frequencies occur early in a stimulus.

**Figure 2.**
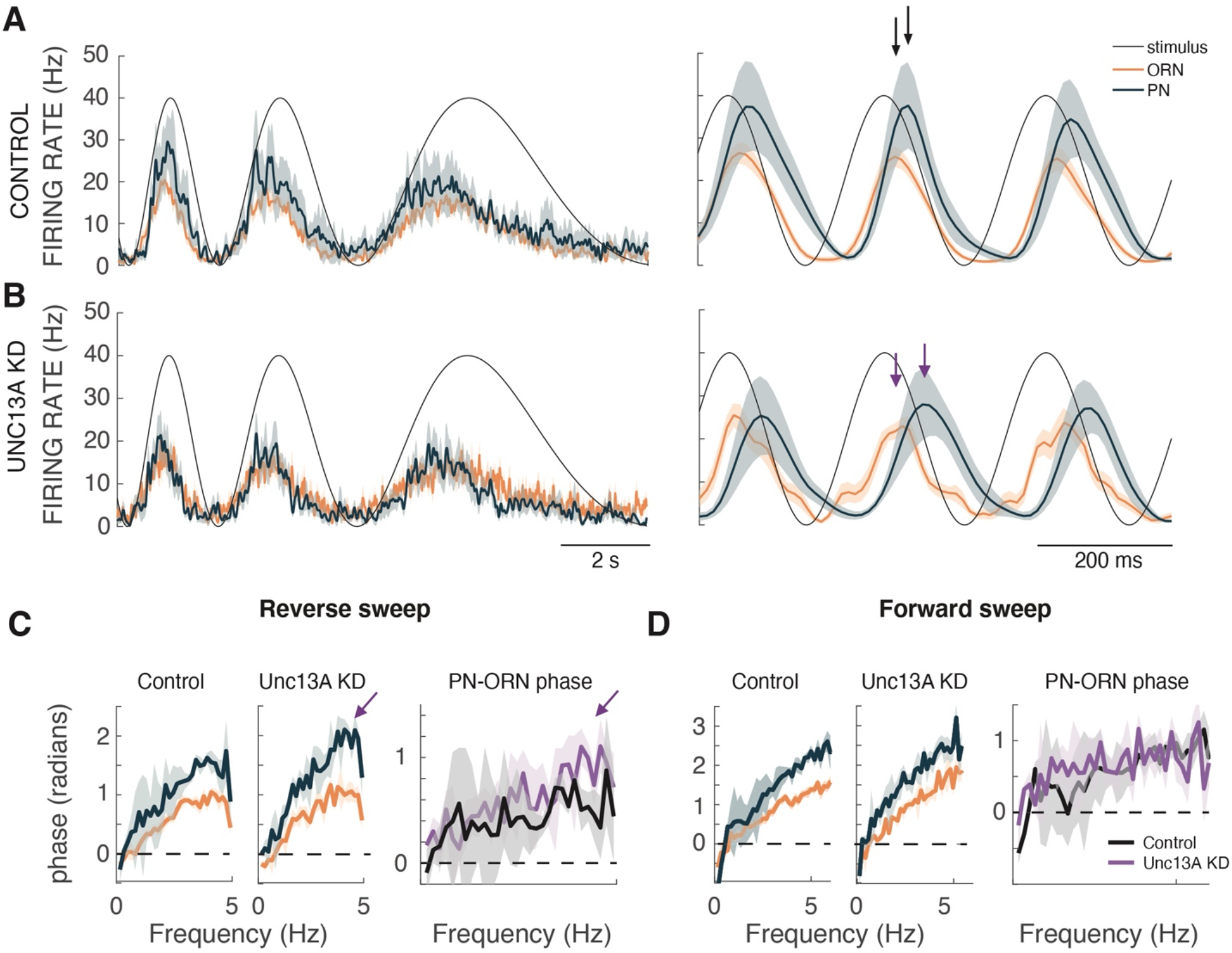
Knockdown of unc13A at ORN-PN synapses increases the latency of PN responses at high stimulus frequencies. (A) Average firing-rate responses of ORNs (orange) and PNs (blue) to a low frequency (left) and high frequency (right) segment of the reverse frequency sweep in control flies (n = 13 ORNs, n = 9 PNs. Mean +/– SEM of resampled jackknife traces. Same data as in Fig. 2 at higher magnification). Firing rate peaks slightly precedes stimulus peak at the lowest frequency and lag the stimulus peak at high frequency. (B) Same as B for unc13A KD (n= 15 ORNs, n = 8 PNs. Same data as in Fig. 2 at higher magnification.) Vertical arrows indicate longer latency to PN peak at high frequency in unc13A KD flies. (C) Left: Response phase (latency to half-max) as a function of stimulus frequency for the reverse sweep. ORN in orange and PN in blue (mean +/– SEM). Right: phase difference between PN and ORN responses for control (black) and unc13AKD (violet). Arrows indicate increased delay in PN responses at high frequencies. (D) Same as C for the forward sweep.

Finally, we examined the effects of unc13A knock down on synaptic gain. We stimulated orco+ ORNs with a series of 2 s light pulses (0 to 120 uW/cm^2^ intensity), both before and after a 10 s long adapting pulse at 60 uW/cm^2^. In control animals, PN responses were generally larger than ORN responses, and PN response gain was reduced following the long adapting pulse (Fig. 3 A,C,E) indicating that slow timescale adaptation arises at this synapse (Nagel et al. 2015, Martelli and Fiala, 2019). Responses in unc13A flies were very slightly reduced compared to control, and PN response gain still dropped after the adapting pulse, indicating that this form of adaptation does not require unc13A (Fig. 3 B,D,F). These results indicate that unc13A makes a small contribution to synaptic gain.

**Figure 3.**
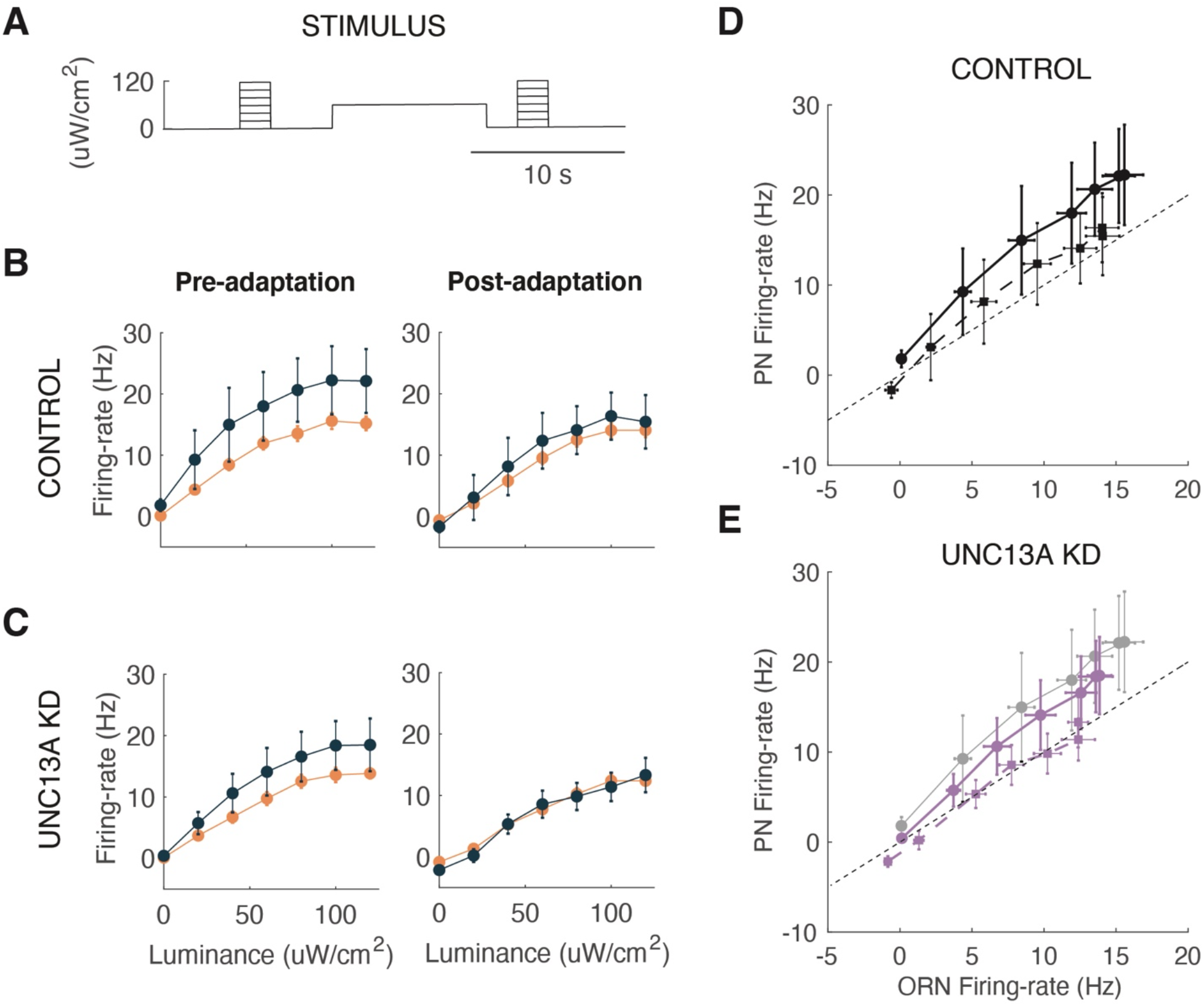
Knockdown of unc13A at ORN-PN synapses reduces the sensitivity of transmission but does not alter slow-timescale adaptation. (A) Stimulus timecourse. Stimulus consisted of a 2 sec test pulse (0 to 120 uW/cm^2^ in 20 uW/cm^2^ intervals) followed by a 10 sec adapting pulse (60 uW/cm^2^), and then a second test pulse at the same intensity as the first pulse. (B) Average firing-rate responses of ORNs (orange; mean +/– SEM; n = 13) and PNs (blue; mean +/– SEM; n = 9) in control flies to the first and second set of stimulus pulses. Light (C) Average firing-rate responses of ORNs (orange; mean +/– SEM; n = 15) and PNs (blue; mean +/– SEM; n = 8) in unc13A KD flies. Same stimuli as in (B). (D) The average PN response is plotted against the average ORN response for control flies (black circle with solid line for baseline; black square with dashed line for adapted). All statistics (mean +/– SEM are calculated using jackknife resampled error). (E) The average PN response is plotted against the average ORN response for unc13A KD flies (violet circle with solid line or baseline; violet square with dashed line for adapted). Statistics as in (E). Control baseline response is replotted in gray for comparison.

Our experiments with frequency sweeps and pulses indicate that unc13A selectively amplifies and accelerates responses to high-frequency stimulus fluctuations with only modest effects on synaptic gain. To examine the effects of unc13 KD on a more natural stimulus, we presented an optogenetic “plume-walk” (Fig. 4A): a stimulus created by taking an upwind trajectory through a ground-level plume movie at fly pace (6mm/s, Connor et al. 2018). Comparing ORN and PN responses to this stimulus in control and unc13A knock-down flies, we found that responses to small plume fluctuations, as would be encountered distant from the odor source, were stronger and more robust in control flies (Fig. 4B,C,D, compare black and purple arrows). Thus, unc13A is likely to play an ethological role in allowing flies to detect odor filaments while far away from an odor source.

**Figure 4.**
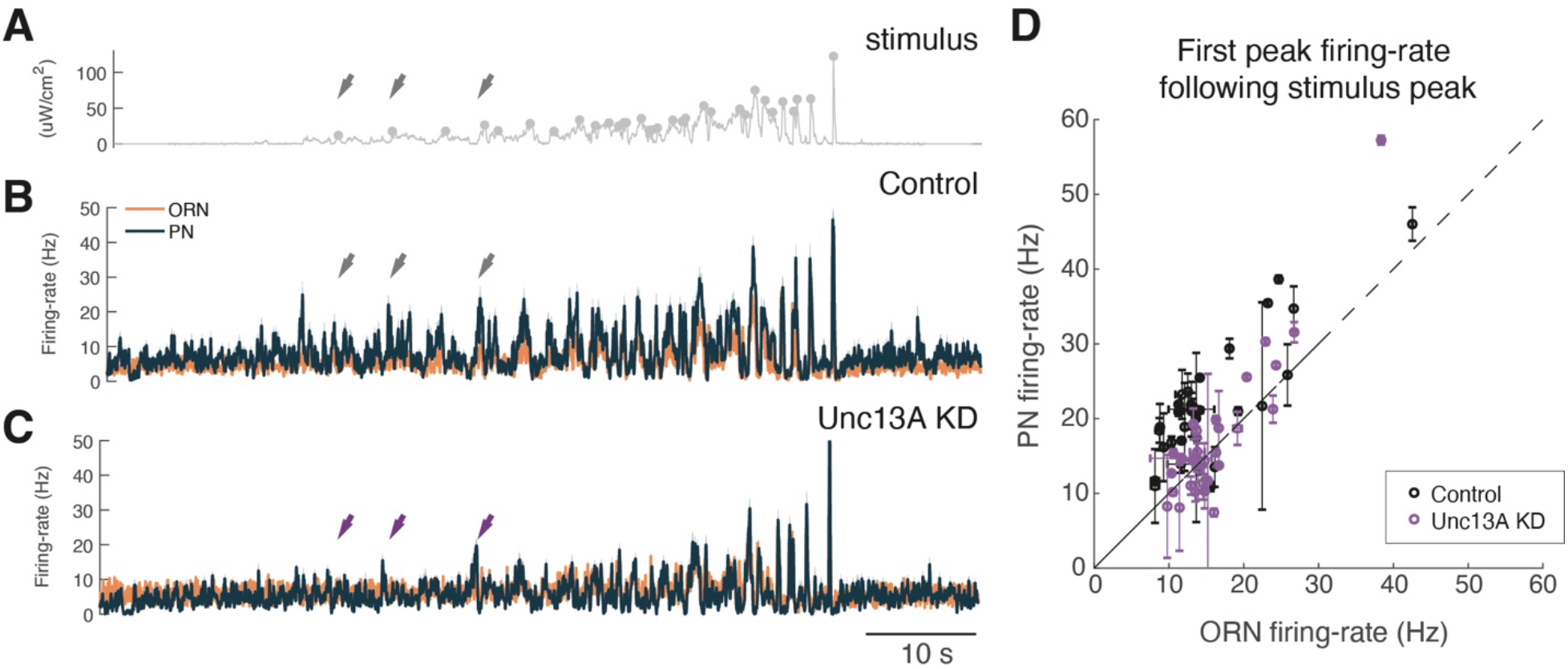
unc13A at ORN-PN synapses amplifies small amplitude ORN inputs during realistic plume stimuli. (A) A virtual “plume-walk” stimulus was constructed by taking an upwind trajectory at fly pace (6 mm/s) through a boundary layer plume measured using planar laser imaging fluorescence and converted to light intensity (Connor and Crimaldi 2018). Luminance values ranged from 0 to 120 uW/cm^2^. Circles denote the stimulus peaks that were used for analysis. (B) Firing-rate responses of ORNs (orange, n=13) and PNs (blue, n=9) to the plume-walk stimulus in control flies. Traces are mean +/– SEM of jackknife resampled traces. Same cells as in Figs. 1–3. (C) Same as B for unc13A KD (n=15 ORNs, n=8 PNs). (D) The average PN response is plotted against the average ORN response for the first peak following each stimulus peak (denoted by a circles in A) for control (black) and unc13A KD (violet). Error bars are standard error of jackknife resampled traces (see methods). Points above the unity line indicate amplification of the response in PNs relative to ORNs. Control flies show greater amplification at most points, especially far from the odor source.

### A two-component model of synaptic transmission can qualitatively replicate many of the observed effects of unc13A knock-down

In a previous study, we developed a two-component model of ORN-PN synaptic transmission, in which a strong rapidly-adapting component, and a weak slowly-adapting component sum to generate total synaptic conductance (Nagel et al. 2015). Knock-down experiments indicate that unc13A contributes only to the fast component of transmission (Fulterer et al. 2018). We therefore asked whether reducing the conductance of the fast component in our model can recapitulate the effects of unc13A knockdown on temporal filtering, phase, and gain.

To address this question, we combined our previous model of synaptic transmission with a GLM model of the transformation from membrane potential to spiking, parameterized by a stimulus filter and a postspike filter (Figure 5A), allowing us to simulate PN responses to diverse ORN responses (Figure 5B–D). We next asked how changing a single parameter of the model— the conductance of the fast component— affects temporal filtering, phase, and gain seen in PN firing-rate responses. Because this manipulation also changed the baseline membrane potential of the model, we added additional offset current (similar to our experiments) to make the resting membrane potential similar between models with and without the fast component of transmission.

**Figure 5.**
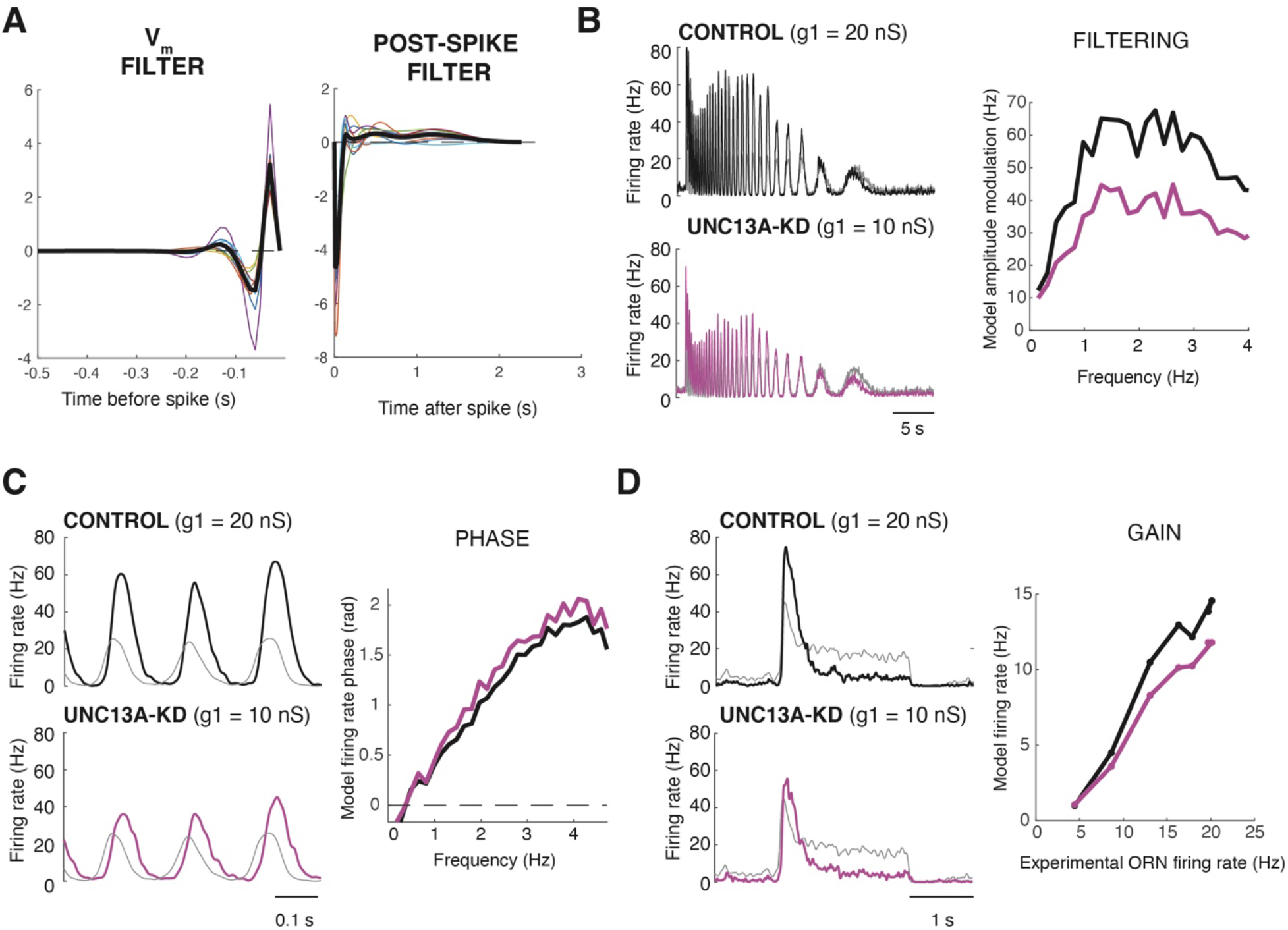
A reduction in the conductance of fast synaptic transmission can account for unc13A-KD-mediated decreases in high-frequency transmission, response latency, and gain. (A) GLM model of the relationship between membrane potential and neural spiking, estimated from PN data. Left: Membrane potential filter. Right: Post-spike filter. See Methods for details. (B) Model PN responses (thick lines) computed from ORN data (thin gray lines) in response to the reverse frequency sweep. Top: model simulating control PNs (black) with g1 (conductance of fast synaptic component) of 20 nS. Bottom: model simulating unc13KD PNs (magenta) with g1=10nS. Right: Reducing the fast synaptic conductance (g1) preferentially reduces the amplitude of responses to higher-frequency inputs. (Modulation depth computed as in Figure 1). (C) Same as (B) but at higher temporal resolution to show phase relationship between ORN data and PN model output at 3 high frequency cycles. Right: Differences in phase between stimulus and PN response normalized by stimulus cycle frequency and plotted as a function of stimulus frequency. Reducing the fast synaptic component (g1) increases the delay of PN firing-rate relative to the stimulus. (D) Model PN responses (thick lines) computed from ORN data (thin gray lines) in response to a light pulse. Colors as in B and C. Right: Reducing the fast component of the synaptic model (g1) from 20 nS to 10 nS reduces the gain of transmission. Mean firing rate as a function of ORN input computed as in Figure 1.

Consistent with our data, we found that reducing the conductance of the fast component from 20nS to 10nS per spike preferentially decreased the response to high frequency ORN firing rate fluctuations versus low frequency firing rate fluctuations (Figure 5B). This computational manipulation also increased the phase of model PN responses relative to ORN responses (Figure 5C). Finally, reducing the conductance of the fast component modestly reduced ORN-PN gain (Figure 5D). This model supports the idea that transmission at ORN-PN synapses arises from two kinetically distinct components, and that the fast component synapses is mediated by unc13A.

### Upwind navigation in response to high frequency stimulus fluctuations depends on unc13A

Olfactory navigation behaviors critically depend on the temporal dynamics of odor input (Schulze et al. 2015, Alvarez-Salvado et al. 2018). In particular, airborne odor plumes generate intermittent high frequency fluctuations in odor concentration, particularly near the plume midline (Celani et al. 2014, Connor and Crimaldi, 2018), and high frequency odor encounters were recently shown to preferentially promote upwind orientation, a key component of odor attraction, in walking *Drosophila* (Demir et al. 2020). We therefore asked whether unc13A promotes upwind navigation in response to high frequency stimulus fluctuations.

To measure behavioral responses to odor, we employed a miniature wind-tunnel apparatus that we had previously designed to make quantitative measurements of olfactory behavior (Alvarez-Salvado et al. 2018). In this paradigm, flies run upwind in response to a pulse of attractive odor (apple cider vinegar), showing increased upwind velocity and groundspeed. Following odor offset, they perform a local search characterized by increased angular velocity. Here we replaced the odor stimulus with optogenetic activation of ORNs, using red LEDs positioned above the arena. We observed upwind running during light, and search behavior at light offset, similar to that observed with odor, over arrange of stimulus intensities (Fig, 6A,B, Matheson et al. 2021).

**Figure 6.**
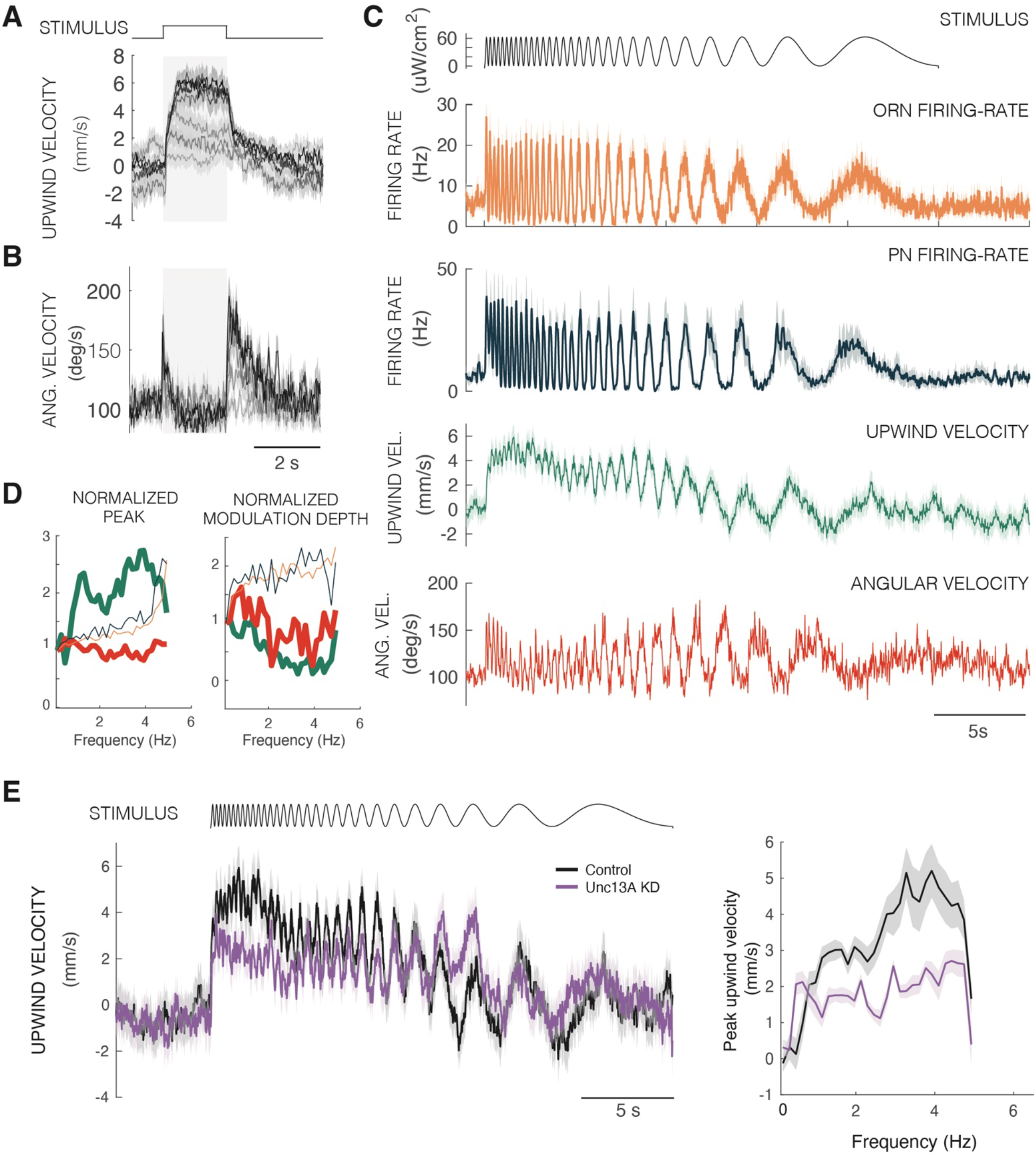
Preferential upwind running in response to high-frequency stimulus fluctuations requires unc13A. (A) Average upwind velocity response across flies (+/– SEM; n = 44 flies) expressing Chrimson under orco-GAL4 to 2 s light pulses from 0 to 120 uW/cm^2^ in 20 uW/cm^2^ intervals. Gray boxes indicate stimulus period. (B) Average angular velocity responses of the same flies to the same stimuli. (C) Neural and behavioral responses to the reverse frequency sweep stimulus shown in Figure ORN data (n= 11) are from heterozygous flies matching behavior data. PN data are from homozygous flies, reproduced from Figure 2 (see Methods). Behavioral responses are mean +/– SEM (n = 44 flies). (D) Peak value and modulation depth (difference between maximum and minimum within each stimulus cycle) plotted as a function of stimulus frequency for each neural response or behavioral parameter. Colors as in (C). All traces are mean +/– SEM of jackknife resampled values. (E) Right: Upwind velocity response to the reverse frequency sweep stimulus for control (black; n = 44) and unc13A KD (violet; n = 40) flies. All traces are mean +/– sem on resampled traces. Left: Average peak upwind velocity plotted as a function of stimulus frequency for each stimulus (mean +/– SEM).

To examine how behavior depends on temporal frequency, we exposed flies to reverse frequency sweep stimuli identical to those used for physiology. In response to this stimulus, we found that the peak upwind velocity increased with stimulus frequency, as seen in ORN and PN responses (Figure 6C). However, modulation depth decreased with stimulus frequency (Figure 6D), leading to a more consistent upwind velocity over time. These data indicate that rapid fluctuations in an odor stimulus— as encoded by high frequency fluctuations in ORN firing rate— can preferentially promote strong upwind orientation. We then performed the same experiment in flies where unc13A was knocked down in ORNs (Figure 6E). In these flies, upwind velocity in response to the highest frequency fluctuations was reduced, and upwind velocity was more constant as a function of stimulus frequency. These data argue that the reduction in high frequency transmission observed between ORNs and PNs is relevant for behavior, and that unc13A plays a role in promoting upwind navigation in response to high frequency odor fluctuations.

## Discussion

### A molecular mechanism for temporal filtering at presynaptic terminals

The frequency content of natural stimuli often contains important information. For example, both the carrier frequency and pulse frequency of song are important for recognizing conspecifics in frogs, birds, and insects (Vignal and Kelley, 2007; Nagel et al. 2010; Coen et al. 2014). Visual motion is a critical cue both for locating prey and for measuring one's own motion through space (Krapp and Hengstenberg, 1996; Bianco et al., 2011); it is detected by comparing signals from nearby points in space that have been differentially filtered in time (Gruntman et al. 2019; Kim et al. 2014). In an odor plume, the frequency of odor encounters increases as an animal approaches the plume midline and the source of the odor, providing a critical cue about the location of food and mates (Celani et al. 2014; Connor et al. 2018). Thus, mechanisms to differentially filter sensory signals in time are critical for proper processing and responses to natural stimuli of all modalities.

One potential mechanism for implementing temporal filters within the brain is short-term synaptic plasticity. When a chemical synapse is activated repeatedly, the postsynaptic response typically either shrinks or grows over time (Abbott and Regehr, 2004). Computationally, short-term depression implements a low-pass filter on spike rate— low-frequency spikes produce large post-synaptic responses, while high-frequency spikes lead to depression. However, when information is carried in neural firing rates, short-term depression may be thought of as implementing a high-pass filter. A depressing synapse transmits a large signal when the presynaptic firing rate changes rapidly, but attenuates fluctuations when they are slow (Abbott et al. 1997). The computational properties of short-term plasticity have frequently been evoked to explain sensory responses observed within the brain (Espejo et al. 2019; Seay et al. 2020). However, it has been challenging to directly test the role of synaptic depression in shaping sensory processing in intact organisms.

At a molecular level, short-term plasticity depends on a host of pre-synaptic and post-synaptic specializations that can be characteristic of particular synapse types (Zucker and Regehr, 2002). On the presynaptic side, depression often arises from depletion of synaptic vesicle pools positioned at different distances from release machinery. In *Drosophila*, two unc13 isoforms—A and B— are located at characteristic nanometer distances from voltage-gated calcium channels, with the A isoform located closest to the calcium channel (Böhme et al. 2016). Knock-down of the A isoform reduces the amplitude of unitary EPSCs at the neuromuscular junction and at ORN-PN synapses, and shifts these synapses from depression to facilitation (Fulterer et al. 2018). Different synapses express different levels of A and B isoforms, and this expression appears to correlate with synaptic dynamics (Fulterer et al. 2018, Pooryasin et al. 2021). Unc13 isoforms that differentially regulate short-term plasticity have also been identified in nematodes (Hu et al. 2013) and in vertebrates (Rosenmund et al. 2002).

Here we took advantage of the genetic control possible in *Drosophila* to directly test the role of short-term plasticity in sensory processing and behavioral responses to dynamic stimuli. We found that knock-down of unc13A specifically at ORN terminals reduced the ability of post-synaptic neurons to follow rapid changes in presynaptic firing rate with short latency. In the context of a natural plume stimulus, unc13A knock-down reduced the ability of the olfactory system to encode distant, small-amplitude plume filaments. Behaviorally, unc13A knock-down reduced upwind running in response to rapid stimulus fluctuations. Thus, high-levels of unc13A expression at ORN terminals are required for this synapse to faithfully transmit the wide range of stimulus frequencies present in natural olfactory signals, and this transmission is critical for proper behavioral responses to rapidly fluctuating odor plumes.

### Contribution of ORN-PN synapses to temporal coding

Because of its genetic and electrophysiological accessibility, the synapse between olfactory receptor neurons and projection neurons has emerged as model for understanding the contributions of synaptic processing to sensory coding (Kazama and Wilson 2008, Kim et al. 2015, Nagel et al. 2015, Martelli and Fiala 2019, Brandão et al. 2021). Work over the past decade has identified several transformations of sensory information that occur at this synapse. First, synaptic transmission amplifies ORN responses, allowing for detection of weak odor signals (Kazama and Wilson 2008). Second, a slow form of adaptation— likely arising from presynaptic depression— originates at this synapse (Nagel et al. 2015, Martelli and Fiala 2019). Third, two kinetically distinct components of synaptic transmission are required to account for PN responses to complex odor dynamics (Nagel et al. 2015). A model incorporating these two components, along with presynaptic inhibition, was previously shown to capture PN responses to dynamic odor stimuli (Nagel et al. 2015).

Here we used a genetic manipulation of ORN-PN synapses, coupled with optogenetic activation of ORNs, to directly link the fast component of synaptic transmission to unc13A, and to show that this fast component is required for proper upwind runs in response to a high-frequency odor stimulus. Together with previous data showing that the fast component could be blocked by the nicotinic acetylcholine receptor antagonist curare (Nagel et al. 2015), these data support the idea that two separable molecular specializations underlie the ability of this synapse to transmit high frequency and low frequency information. Further, they argue that both pre- and post-synaptic specializations are required for each frequency component. In the future it will be interesting to investigate whether there is coordination between the expression of presynaptic and postsynaptic components that tune a synapse for low- or high-frequency transmission.

### *Drosophila* navigation as a model to link molecular deficits to behavioral phenotypes

A major goal of neuroscience research is to understand the relationship between synaptic molecular deficits and behavioral phenotypes. One challenge in linking these two is that the timescales at which neural and synaptic activity are measured (milliseconds to seconds) are typically much faster than the timescales at which behavior is measured. Navigation behaviors in *Drosophila* present an attractive model for relating synaptic deficits to behavior. Navigation provides a rich real-time analog read-out of dynamic sensory input (Schulze et al. 2015, Alvarez-Salvado et al. 2018, Demir et al. 2020). Behavior can be driven by optogenetic activation of sensory inputs, allowing for precise temporal control of sensory input (Schulze et al. 2015). Further, navigation behaviors can be modulated on short timescales by activation of dopaminergic neurons (Handler et al. 2019). The wealth of genetic tools for manipulating synaptic function, coupled with the ability to easily target specific molecular manipulations to specific synapses, should allow for a precise biophysical dissection of how a dynamic stimulus is transformed into a dynamic behavior, and how synaptic deficits at different levels of processing differentially impact behavior.

One advantage of using navigation as a behavioral model is that different components of the behavior depend on different temporal aspects of the stimulus. For example, upwind orientation is promoted by high-frequency fluctuations, however, initiation of movement depends on an integration of odor information across multiple encounters (Demir et al. 2020). In general, the time course of behavior was much slower than the time course of PN firing, suggesting that some type of low-pass filter is implemented between PN firing and the generation of behavior. Identifying where and how each of these temporal filters arises in the fly nervous system can provide insight into how brains implement one of the basic computational components of signal processing.

## Acknowledgements

The authors would like to thank Stephan Sigrist for generously sharing unc13A RNAi flies and Nick Stavropoulos for w1118 5905 flies. Stephan Sigrist and Jonathan Victor as well as members of the Nagel and Schoppik labs provided helpful comments on the manuscript. This work was funded by MH109690 to K.I.N.

## Methods

### Fly stocks

Fly strains were reared under standard laboratory conditions and a 12/12-hour light/dark cycle at 25°C.

To perform optogenetic experiments we generated a stable stock norpA;orco-Gal4;UAS-Chrimson that allowed us to activate orco+ ORNs using red light in a genetically blind background. The norpA transgene had been backcrossed seven generations to an isogenic w1118 wild-type selected for their robust walking behaviors (Bloomington #5905, referred to elsewhere as iso31, Ryder et al., 2004) and was previously used for behavioral experiments in Alvarez-Salvado et al. 2018.

To knock down unc13A expression we used an RNAi construct against unc13A developed by the Sigrist lab (Fulterer et al. 2018). We backcrossed this stock 5 generations into the w1118 5905 line.

For control ORN and PN recordings we used homozygous flies of the genotype norpA;orco-Gal4;UAS-Chrimson. For unc13A KD ORN and PN recordings we used hemizygous males norpA/y;orco-Gal4/+;UAS-Chrimson/unc13A RNAi. In all figures we directly compare PN recordings to ORN recordings from the same genotype.

For control behavioral recordings we crossed the norpA;orco-Gal4;UAS-Chrimson line to 5905 wildtype flies and used hemizygous males norpA/y;orco-Gal4/+;UAS-Chrimson/+. To compare behavior to ORN recordings, we also recorded ORN responses in these hemizygotes. ORN data in Figures 6 is from genotype matched flies, while PN data is from homozygotes (see figure legends). Unc13A KD behavior was also performed in hemizygote males norpA/y;orco-Gal4/+;UAS-Chrimson/unc13A RNAi.

For ORN and PN recordings, experiments were performed 3-day post eclosion. Adult flies were placed on agar with all trans-retinal (ATR) for 48 – 72 hours prior to recordings.

For behavior, adult flies were placed on agar with all trans-retinal (ATR) for 48 hours and at room temp in custom-made boxes with a 12/12 hr light dark cycle. For 24 hours prior to the experiment, the flies were starved by placing them in a vial containing only KimWipes soaked with distilled water to humidify the air. Experiments were performed on flies that were approximately 3-5 days old and between 2-4 hours after the lights came on (ZT2 – ZT4).

**Key Resources Table.**

| Reagent/resource | source | Identifier |
| --- | --- | --- |
| w- *NorpA* (5905) | Stavropoulos lab |  |
| orco-gal4 | Vosshall Lab; Wang et al (2003) |  |
| 20x-UAS-IVS-csChrimson | BDRC | RRID: 55136 |
| unc13 isoA RNAi<br>backcrossed 5 generations<br>into 5905 | Sigrist Lab<br>Backcrossing performed for<br>this study |  |
| Software and algorithms |  |  |
| Matlab | Mathworks | R2017b |
| Python |  |  |

### ORN recordings

Extracellular sensillum recordings from ORNs on the antenna were made as described in Nagel and Wilson, 2011. We targeted large basiconic sensilla, each of which contains 2-4 ORNs (de Bruyne, 2001). Electrical signals were acquired using a Model 2400 amplifier (A-M Systems) and low-pass filtered at 2 kHz with a Brownlee Precision 410 amplifier before digitization at 10 kHz using a National Instruments PCI-6321 card. Spikes from different ORNs were separated based on amplitude using custom Matlab scripts.

### Whole-cell patch clamp recording

*In vivo* whole-cell patch clamp recordings from PNs were performed as described in Nagel et al. 2015. PNs were identified by their characteristic location and size within the antennal lobe. One PN was recorded per brain. Patch-clamp electrodes (6 – 8 MⅢ) were filled with potassium-aspartate intracellular solution (Wilson and Laurent, 2005), which contained 140 mM of potassium hydroxide, 140 mM of aspartic acid, 10 mM of HEPES, 1 mM of EGTA, 1 mM of potassium chloride, 4 mM of magnesium adenosine triphosphate, and 0.5 mM of trisodium guanine triphosphate, and 13 mM of biocytin hydrazide. A small constant hyperpolarizing current was injected during PN recordings, beginning immediately after break-in, in order to bring the membrane potential between –50 to –60 mV. Voltage was acquired in current-clamp mode using a Model 2400 amplifier (A-M Systems) at 10 kHz. Spikes were extracted using custom MATLAB scripts that filtered, differentiated, and thresholded the raw voltage traces.

### Behavioral apparatus

Behavioral measurements were made using a miniature wind tunnel apparatus described previously (Alvarez-Salvado, 2018). Flies were constrained to walk within a shallow rectangular arena. A constant airflow traveled through the arena at 12 cm/s as measured by a calibrated hot wire anemometer (Dantec Dynamics MiniCTA 54T42). An LED panel consisting of interleaved strips of IR and red LED strips (environmental lights) was placed over the arena. Light stimuli as described above were randomly interleaved and delivered every 30 s.

Analysis and filtering of behavioral data was performed according to methods described in Alvarez-Salvado et al., 2018. We tracked the position and orientation of flies in real time using custom Labview software (National Instruments) and further analyzed the data in MATLAB (Mathworks). We discarded any trials with tracking errors and trials in which flies moved less than 25 mm. Behavioral parameters (ground speed, upwind velocity and absolute angular velocity) were computed only using segments in which flies moved at speeds greater than 1 mm/s. We additionally excluded data after the fly reached the upwind end of the chamber before the end of the trial.

### Stimulus delivery and calibration

For electrophysiological recordings, stimuli were delivered using a single red LED (628 nm, environmental lights) mounted on a micromanipulator approximately 25cm from the fly. For behavioral experiments, red light was delivered through a set of red LED strips (628 nm, environmental lights) mounted above the wind tunnel arenas, and interleaved with the IR LEDs used for visualizing the flies.

Light power for both electrophysiology and behavior was controlled through Lux drive LED buck drivers (3021/3023 BuckPuck; Luxdrive LED Dynamics) using an analog output voltage signal through a National Instruments data acquisition board (PCIe-6231). The voltage was adjusted to obtain the desired light intensities over the linear dynamic range of the buck puck as measured with a photometer (S120VC, ThorLabs) placed at the location of the fly. The light intensities used were selected from a preliminary set of experiments measuring PN firing-rate responses to linear increments of light power. A set of seven intensities ranging from 0 to 120 uW/cm^2^ in 20 uW/cm^2^ increments was found to traverse the dynamic range including saturation of PN firing-rate responses and without inducing depolarization-block. We therefore decided to use these light intensities for all recordings including extracellular sensillum recordings of ORNs, whole-cell patch clamp of PNs, and behavior.

To measure intensity coding and adaptation, we generated a stimulus consisting of two test pulses (0- 120 μW/cm^2^ in increments of 20μW/cm^2^), each 2 s in duration, separated by a 10 s adapting step at 60 μW/cm^2^; these steps were separated by 2 s. To measure frequency and phase encoding, we generated frequency sweep stimuli that smoothly transitioned through frequencies 0.1 to 5 Hz in both forward and reverse directions and through intensities 0 – 60 μW/cm^2^. We also constructed a ‘plume walk’ stimulus by taking an upwind trajectory at fly pace (6 mm/s) through a boundary layer plume measured using planar laser imaging fluorescence (Connor et al., 2018).

### Analysis of neural responses

All means and standard errors of firing rate data – for plotting average traces and summary statistics - were found by resampling. Jackknife resampled traces were calculated by calculating the mean of n-1 trials and repeating this process n times, where n is the number of trials. The jackknife estimate for the mean was taken as the mean of the jackknife estimates and the jackknife estimate of the standard deviation of the sample mean was taken as the standard deviation of the jackknife estimates of the sample mean, which was given as:

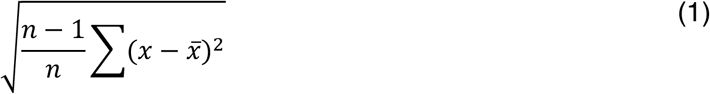

Frequency responses in Figures 1 and 6 were calculated on jackknife resampled traces. Modulation depths were taken as the difference between minimum and maximum responses in each stimulus chirp cycle of each resampled trace. The standard error in each stimulus chirp cycle was given by equation (1). The frequency of the cycles was estimated as 1 over the duration of the cycle. The standard error at each time point is given by equation (1).

ORN and PN phase were computed by calculating the latency to half-maximum on each cycle. This was calculated from jackknife resampled traces and normalized by 2πf. Mean and standard error was obtained as in equation 1.

Average responses for pulses in Figures 3 were calculated by taking the average of jackknife estimates for each time point. The average was taken over the entire 2 s stimulus interval. All statistics were extracted from baseline-subtracted traces, where baseline was taken as the 2 s preceding the stimulus.

Peak responses during the plumewalk were calculated by first detecting maxima in the plume stimulus and then finding the first peak in the ORN and PN jackknife resampled traces immediately following each stimulus peak.

### Synaptic depression model

We modeled PN firing-rate responses to ORN firing-rate inputs using a combination of a biophysical model of a two-component synapse (Nagel et al. 2015) and passive membrane with a GLM that translates model PN membrane potential into firing-rate (Weber & Pillow, 2017).

To model short-term depression in the ORN-PN synapse, we used a classical model of short-term plasticity (Abbott et al., 1997; Nagel & Wilson, 2015).

The input to the model was measured ORN firing-rate. The ORN firing-rate produced a change in the postsynaptic conductance with two components, fast and slow. Each component of the synaptic conductance was described by a pair of equations:

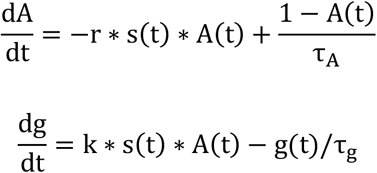

A(t) governs the amplitude of the postsynaptic conductance, s(t) is the average experimentally measured ORN firing rate with units of spikes/ms, r governs the depression rate and τ_A_ is the rate of recovery from depression. The conductance is g(t), where k controls the amplitude of the conductance and τ_g_ is the conductance decay rate. For the fast component, we used r = 0.23 per spike, k = 20 nS per spike, and τ_G_ = 9.3 ms. For the slow component, we used r = 0.0073 per spike, τ_A_ = 33247 ms, k = 1.8 nS per spike, and τ_g_ = 80 ms.

The fast and slow conductances arising from ORN firing-rate were summed to produce a total excitatory postsynaptic conductance in a passive membrane model PN:

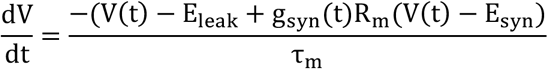

Where V(t) is the membrane potential, E_leak_ is the leak reversal potential, R_m_is the membrane resistance, and τ_m_is the membrane time constant. Parameters were taken from Nagel & Wilson, 2015. To model the effects of unc13A knock-down on temporal filtering, phase, and gain of PN spiking responses we set the conductance of the fast component to 10nS instead of 20nS. In experiments, we injected current to offset a change in conductance such that both the control (g1=20 nS) and the unc13A knock-down (g1=10 nS) conditions the membrane potential sat at ~−50 mV.

To model PN spiking, we fit a GLM using low-pass filtered PN membrane potential as input and spiking as output to extract a membrane potential filter and a post-spike filter (see next section section). Using these filters and the model membrane potential, we simulated GLM spikes in 1 ms time bins and calculated the spike-count probability in a given time bin as a Bernoulli process, a discrete-time version of the Poisson process:

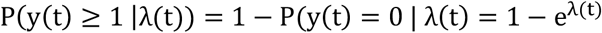

Where λ(t) is the average firing-rate in time bin t.

The PSTH smoothed with a 50 ms was calculated over all spike counts to generate a spike-rate in response to model membrane potential.

### Fitting temporal filters to spike-train data

To model the relationship between membrane potential and spiking we fit Poisson generalized linear models (Weber & Pillow, 2017). We used low-pass filtered PN membrane potential as input to obtain a membrane potential filter, which describes how spike-generating mechanisms integrate Vm. We also fit a post-spike filter, which captures the influence of spike history on the probability of spiking.

To fit the filters, we fit coefficients on a raised cosine basis to reduce the dimensionality and to ensure smoothness of the filters.

The basis vectors were of the form:

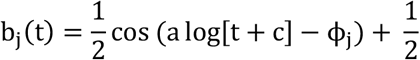

For t such that a log[t + c] ∈ [ϕ_j_ − π, ϕ_j_ − π] and 0 elsewhere. The parameter c determines the spacing of the peaks, with smaller values producing more nonlinearly-spaced peaks. For the stimulus-filter, we used 10 basis vectors to fit a 10 s stimulus filter and 7 basis vectors to fit a 250 ms post-spike filter. For the membrane potential filter, most of the structure was in the first 50-100 ms. In general, we tried to use as few basis vectors as possible with appropriate spacing to represent the entire space. In total, we estimated the coefficients on 17 basis vectors and a DC bias parameter.

For neural spiking, we found the weights on the basis vectors by maximizing the log-likelihood subject to a ridge penalty. In particular, we minimize the posterior distribution by minimizing the negative log-likelihood with an added ridge prior to regularize the filter weights:

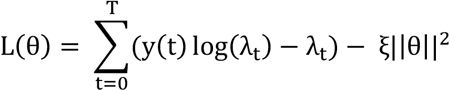

where λ_t_ = X^T^θ_stim_ + Y^T^θ_spike_, θ is a vector representing the weights on the basis functions, y(t) is the spike count at time t, and ξ is a ridge penalty.

For each neuron, we trained on 10 random trials and performed 5-fold cross-validation to choose the value ξ that maximized the likelihood averaged across different cross-validation folds of the data.

We extracted the filter weights by fitting the model to the training data. We calculated the goodness-of-fit using root mean-squared-error by comparing a PSTH smoothed with a 50 ms time-window on GLM simulated spikes and experimental spikes.

